# Expression screening and functional verification of recombinant hirudin from novel chassis cell *Chlamydomonas reinhardtii*

**DOI:** 10.1101/2025.09.25.678210

**Authors:** Li Feng, Junqiao Xing, Aijie Tian, Wei Wang, Yu Zhang, Weihua Wang, Zhangfeng Hu

**Affiliations:** Institute of Microalgae Synthetic Biology and Green Manufacturing, School of Life Sciences, Jianghan University, Wuhan, 430056 Hubei, China; Wuhan Originem Biotech Company Limited, Wuhan, 430056 Hubei, China

**Keywords:** *Chlamydomonas reinhardtii*, synthetic biology, recombinant hirudin, high-density heterotrophic cultivation, anticoagulation

## Abstract

Hirudin is a potent anticoagulant peptide whose clinical application is limited by scarce natural supplies and the suboptimal activity of recombinant versions from conventional microbial hosts lacking key post-translational modifications. Here, for the first time, we establish the “Generally Regarded as Safe” (GRAS) microalga, *Chlamydomonas reinhardtii*, as a novel biological chassis for producing fully functional hirudin. Using high-density heterotrophic fermentation, we generated a bioactive lyophilized algal powder suitable for oral delivery. Crucially, the algal-derived hirudin underwent proper tyrosine sulfation—a key modification absent in prokaryotic hosts—conferring exceptionally high thrombin-inhibitory activity (up to 20,000 ATU/mg). When orally administered to a murine thrombosis model, this hirudin-loaded alga demonstrated potent antithrombotic efficacy. It also exhibited a superior safety profile, showing no signs of the thrombocytopenia or bleeding associated with traditional anticoagulants. This study establishes a green, scalable biomanufacturing and oral delivery platform for therapeutics, highlighting transformative potential at the nexus of agricultural bioengineering, functional foods, and biomedicine.

## Introduction

Microalgae are increasingly recognized as sustainable biofactories for high-value compounds and recombinant proteins, owing to their rapid growth, minimal input requirements, and amenability to genetic engineering^*1-4*^. Among them, *Chlamydomonas reinhardtii* (*C. reinhardtii*) has emerged as a versatile chassis capable of both photoautotrophic growth and high-density heterotrophic fermentation using acetate as a sole carbon source^*5*^. As a unicellular eukaryote, *C. reinhardtii* possesses the machinery for complex post-translational modifications (PTMs), including sulfation, proteolytic processing, and protein folding — while remaining highly tractable to nuclear engineering^*6,7*^. These attributes, combined with its “Generally Regarded as Safe” (GRAS) status^*8*^, make it a prime candidate for developing ingestible functional ingredients and oral therapeutics. This potential for oral delivery is further enhanced by its functional attributes; a polysaccharide-rich cell wall and motile flagella may prolong residence in gastrointestinal fluids and have been leveraged in novel delivery concepts^*9-11*^.

Hirudin, a disulfide-rich anticoagulant peptide from medicinal leeches, prevents thrombosis by binding thrombin with high affinity and blocking the conversion of fibrinogen to fibrin^*12*^. Several hirudin-based drugs and analogs (e.g., desirudin, lepirudin, bivalirudin) have achieved clinical utility across peri-operative and interventional settings for their potent antithrombotic effects. These agents represent a crucial class of therapeutics for managing thromboembolic diseases^*13-16*^.

Despite their efficacy, current hirudin modalities face practical constraints: a short systemic half-life necessitating frequent parenteral administration, increasing patient burden and risk of infection, while their use is associated with class-wide bleeding risks^*17,18*^. Other widely used anticoagulants, such as heparin, carry an additional risk of severe side effects like immune-mediated thrombocytopenia (HIT)^*19-21*^. Therefore, a robust, orally deliverable anticoagulant produced through a sustainable process would hold considerable translational value.

Industrial production of recombinant hirudin has predominantly relied on *E. coli* and yeast^*22,23*^. While cost-effective, these systems lack the machinery for key PTMs — most notably tyrosine sulfation, which is essential for maximal anticoagulant potency^*24*^. In contrast, *C. reinhardtii* possesses eukaryotic PTM pathways, offering a biologically relevant milieu for producing fully active peptides^*6,7,25,26*^. Therefore, this study was designed to harness these attributes to overcome the limitations of current hirudin production and delivery. Our objective was to engineer *C. reinhardtii* to express recombinant hirudin, establish a scalable high-density fermentation process, and evaluate the *in vivo* antithrombotic efficacy and safety of this novel, orally administered microalgae-based therapeutic.

## Materials and methods

### Strains, plasmids, and animals

The wild-type *C. reinhardtii* strain CC-1690 (*21gr*) and the original expression vector pHyg3 were obtained from the Chlamydomonas Resource Center (University of Minnesota, St. Paul, MN, USA) and maintained by routine subculture. C57BL/6J mice were purchased from Beijing Vital River Laboratory Animal Technology Co., Ltd. (Beijing, China). All animal experimental protocols were reviewed and approved by the Animal Ethics Committee of Jianghan University. Mice were housed in a specific-pathogen-free (SPF) facility under a controlled 12-hour light/dark cycle at a constant temperature of 23 ± 2°C and a relative humidity of 30–70%.

### Construction of the recombinant hirudin expression vector

The recombinant expression vector was constructed using the pHyg3 plasmid as a backbone. A core promoter module was assembled by fusing the endogenous *Chlamydomonas TUB2* promoter, the 5′ region of the hygromycin resistance gene (*Hyg*), and the first intron of *rbcS2* gene. Downstream of this module, the hygromycin resistance gene, the FMDV 2A self-cleaving peptide sequence, and a codon-optimized hirudin coding sequence (optimized by website https://www.novopro.cn/tools/codon-optimization.html) were sequentially inserted. An HA tag was added to the C-terminus of the coding sequence to facilitate detection. The fully assembled plasmid was transformed into competent *Escherichia coli* DH5α cells (Vazyme, C502) for amplification and subsequently verified by Sanger sequencing (Sangon Biotech Co., Ltd.). The sequence-verified plasmid was linearized by digestion with the KpnI-HF restriction enzyme (New England Biolabs, R3142S) at 37°C for 30 minutes, followed by recovery of the target fragment using a gel extraction kit (Sangon, B110092).

### *C. reinhardtii* transformation and positive clone screening

*C. reinhardtii* cells were cultured in Tris-Acetate-Phosphate (TAP) medium^*27*^ at 24°C with continuous aeration under a 14 h light/10 h dark cycle at 8000 lux. Cells were grown to the logarithmic phase (approx. 10^7^ cells/mL) and subsequently diluted with fresh TAP medium to a concentration of 4×10^6^ cells/mL. For transformation, cells were collected by centrifugation and mixed with 100–200 ng of linearized plasmid DNA. The mixture was transformed via electroporation using a BTX ECM630 electroporator^*28*^. After a recovery period on a gentle shaker, the cells were plated onto TAP agar containing 25 mg/L hygromycin B (Merck, 400052). The plates were incubated under an inverted light cycle to select for transformants.

### Western blot analysis

Putative transformants were screened by Western blot. Harvested cells were lysed in a chilled buffer supplemented with SDS and a protease inhibitor cocktail (Roche, 04693132001), and the protein samples were denatured by boiling for 10 minutes. The expression and sulfation status of recombinant hirudin were detected using an anti-HA-Peroxidase antibody (Roche, 11867423001) and an anti-sulfotyrosine monoclonal antibody, Sulfo-1C-A2 (Abcam, ab136481), respectively, with Histone H3 as a loading control.

### High-density heterotrophic fermentation and biomass processing

Fermentation of *C. reinhardtii* was conducted in a bioreactor (Baoxing Biotech, BIOTECH-5JG) using Tris-Acetate-Phosphate (TAP) medium as the initial culture medium. The pH was maintained at 7.8 ± 0.2 using 1 M NaOH and 20% (v/v) acetic acid. Corn oil containing 30% (w/w) polyglycerol (Macklin, T909687) was used as an antifoaming agent. Seed cultures, grown in flasks to a density of 10^6^ cells/mL, were used for inoculation. The fermentation was maintained at 26 ± 0.3 °C with an initial agitation of 100 rpm and an aeration rate of 4 L/min. The dissolved oxygen level was kept above 30% saturation. At approximately 48 hours post-inoculation, a feeding strategy was initiated using a 50× Tris-free TAP concentrate supplemented with 0.5 M sodium acetate and 20% (v/v) acetic acid. The feed rate was calculated based on an exponential model^*29*^, as shown in Equation (1):

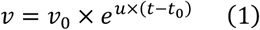

Where *v* represents the fermentation process feeding rate (mL/h), *v*_*t*_is the initial feeding rate (mL/h), *et*is a natural constant, *ut*is the specific microbial growth rate, *tt*is fermentation time, and *tt*_*t*_is the time at which feeding was initiated.

The culture was harvested by centrifugation (4500 rpm) after 144–168 hours, once the cell density reached approximately 5×10^8^ cells/mL. The resulting biomass was lyophilized (Ningbo Xinzhi Biotechnology, ZLGJ-18), then ground and sieved to obtain a fine powder.

### Determination of hirudin anticoagulant activity

The anticoagulant activity of recombinant hirudin was quantified using a colorimetric thrombin neutralization assay kit (GeneMed, GMS10147.2). Following the established definition, one anticoagulant unit (ATU) is the amount of hirudin required to neutralize one International Unit (IU) of thrombin^*30*^. Recombinant hirudin samples produced in *E. coli*, yeast, and *C. reinhardtii* were diluted to a working concentration of 50 μg/mL using Buffer A supplied with the kit. The assay was performed in a 96-well microplate. To each well, 120 μL of Buffer A, 20 μL of Reaction Solution B, 20 μL of Substrate Solution C, and 20 μL of the diluted sample were added sequentially. The mixture was pre-incubated for 1 minute at 25 °C before the addition of 20 μL of color-developing solution. The reaction was then allowed to proceed for 60 minutes at 37 °C in the dark. The absorbance was measured at 405 nm with a microplate reader. A standard curve was generated using a hirudin standard serially diluted to concentrations from 0 to 4 μg/mL. The thrombin inhibition rates and the specific activity of the hirudin samples (ATU/mg) were calculated by interpolating from this standard curve.

### Stability analysis

To evaluate the long-term stability of the recombinant protein, dried biomass was prepared from *C. reinhardtii* strains cultured in both flasks and a bioreactor. For the analysis, 0.5 g aliquots of the resulting algal powder were mechanically compressed into tablets. The tablets were stored for up to 8 months under dark, dry conditions at room temperature (25 ± 0.5 °C). At specified time points (1, 3, 5, 6, and 8 months), samples were collected to evaluate protein integrity. The presence and stability of the recombinant hirudin were qualitatively determined by Western blot, using an anti-HA antibody (Roche, 11867423001) for detection.

### *In vivo* anticoagulant and antithrombotic efficacy assay

All animal experiments were conducted in compliance with institutional guidelines for animal care and use. Mice were acclimated for three days before the start of the study and subsequently treated once daily at 9:00 a.m. for eight consecutive days. The animals were assigned to one of three treatment groups: (1) oral gavage administration of 0.1 mg/kg recombinant hirudin contained within *C. reinhardtii* biomass; (2) intraperitoneal (i.p.) injection of 2 mg/kg bivalirudin (MCE, HY-P1929); or (3) i.p. injection of 4 U/mouse heparin (MCE, HY-17567C). Following the final treatment on the eighth day, blood was collected from the tail vein for platelet analysis using an automatic hematology analyzer (Pukang Electronics, PE-6800VET). Immediately after blood collection, mice were anesthetized to induce thrombosis in the inferior vena cava (IVC) based on a previously established ligation model^*31*^. While maintaining body temperature at 37 °C, a midline laparotomy was performed, the IVC was exposed, and a complete ligation was made below the renal veins using a 7-0 silk suture before closing the incision. Twenty-four hours post-surgery (on day 9), the mice were euthanized. A gross necropsy was performed to assess for any signs of abnormal bleeding, after which the ligated IVC segment was excised and photographed. The thrombus was then carefully isolated from the vessel and its wet weight was recorded to quantify the antithrombotic efficacy of the treatments.

### Data analysis

All enzymatic activity and stability assays were performed in triplicate. The resulting data were analyzed and graphed using GraphPad Prism software (version 8.0.2). All data were presented as mean ± SEM. Values of P < 0.05 were considered statistically significant.

## Results and Analysis

### Expression of recombinant hirudin in algal strains

The recombinant hirudin polypeptide sequence expressed in *C. reinhardtii* (as reported in GenBank: QDZ37419.1) is N-MFSLKLFLVLLAVCICVSQANRYSVCTETGQNLCL CEGSDLCSLDNHCEIGSNGKNRCVKGEGKPKKPQSNSDLPEEKYEPIPIEDYDK-C^*32*^. After codon optimization, the corresponding nucleotide sequence was modified as follows: ATGTTTTCGCTGAAGCTCTTCCTGGTGCTGCTGGCTGTGTGCATTTGCGTGTCGCAAGCCAACCGCTATTCCGTGTGCACGGAGACCGGCCAAAACCTGTGCCTCTGCGAGGGTTCCGACCTGTGCAGCCTGGACAACCACTGCGAGATCGGCTCGAACGGCAAGAACCGGTGCGTCAAGGGTGAGGGGAAGCCCAAGAAGCCCCAAAGCAACTCCGACCTGCCCGAGGAGAAGTACGAGCCCATCCCTATCGAGGACTACGACAAG. Using the pHyg3 vector backbone, we constructed a recombinant expression plasmid harboring the codon-optimized hirudin sequence (Figure 1A). Transformation of *C. reinhardtii* yielded a low frequency of stable integrants (transformation efficiency *t* 0.3%). Western blot screening of transformant colonies with an anti-HA antibody identified a subset of clones as positive for hirudin expression (Figure 1B). Further immunoblotting of HA-positive colonies with an anti-sulfotyrosine antibody detected a clear sulfation signal that was absent in the wild-type control (Figure 1C), indicating that *C. reinhardtii* successfully produced hirudin with the essential tyrosine sulfation modification.

**Figure 1.**
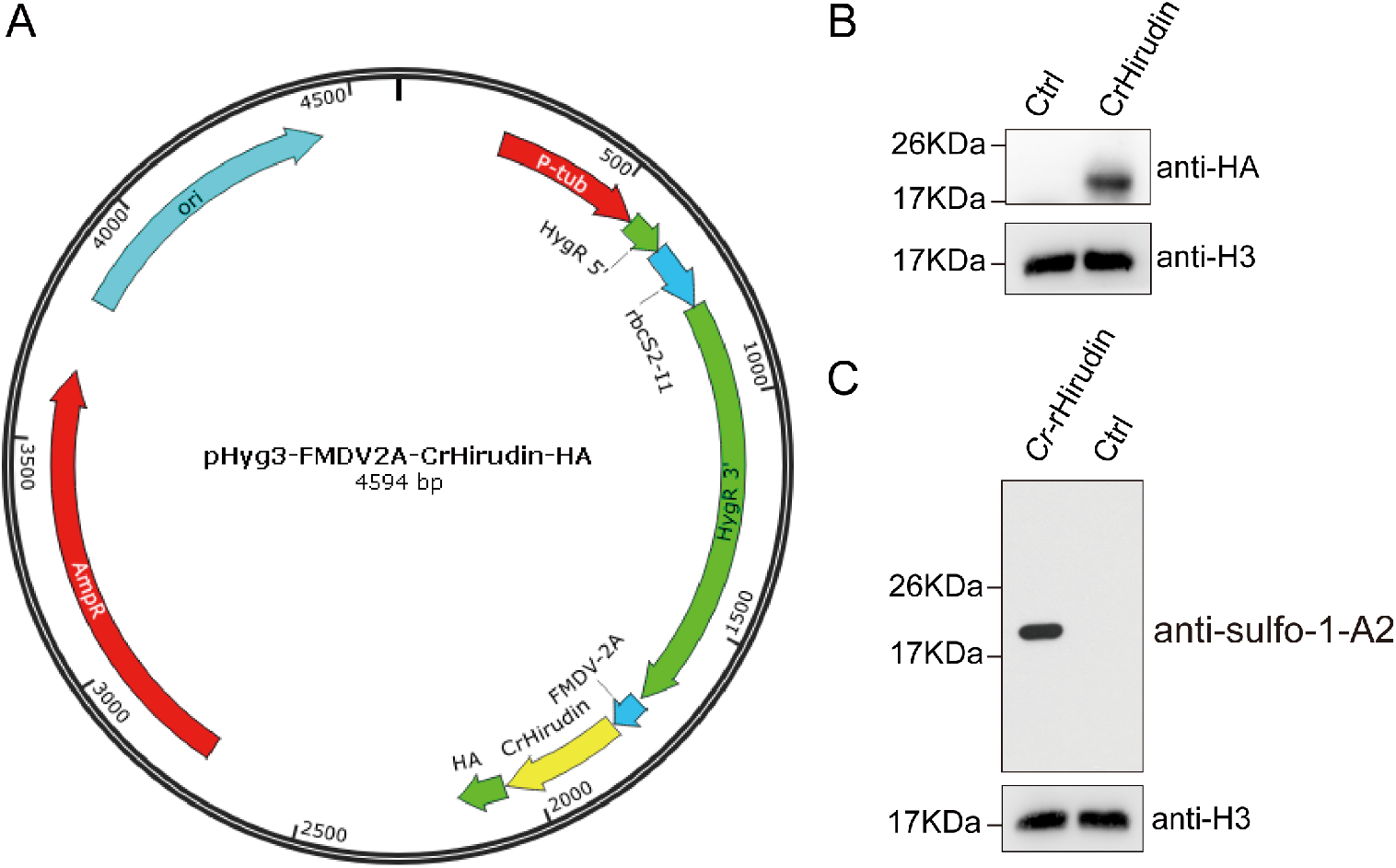
Construction and screening of hirudin-expressing *C. reinhardtii* strains. (A) Schematic of the 4,594 bp expression plasmid, which includes the Chlamydomonas TUB2 promoter, the 5’ region of the hygromycin resistance gene (*Hyg*) with the first intron of *rbcS2*, the FMDV 2A self-cleaving peptide, the codon-optimized hirudin coding sequence, and a C-terminal HA tag. (B) Western blot detection of HA-tagged hirudin in wild-type *21gr* and representative transformant colonies. Histone H3 was used as an internal reference. (C) Western blot using an anti-sulfotyrosine antibody to detect sulfated hirudin. Histone H3 was used as an internal reference.

### Characterization of recombinant hirudin algal powder

Recombinant *C. reinhardtii* expressing hirudin was cultivated using high-density heterotrophic fermentation, and the resulting biomass was processed into a dry powder by lyophilization (Figure 2A). We compared the anticoagulant activity of hirudin produced in different systems – *E. coli, Pichia pastoris* (*P. pastoris*), and *C. reinhardtii* – using a thrombin neutralization assay. Hirudin derived from *C. reinhardtii* exhibited a dramatically higher thrombin-inhibitory activity, reaching approximately 20,000 ATU/mg, whereas the hirudin from *E. coli* and *P. pastoris* showed only a fraction of this activity (Figure 2B). The difference was highly significant (***P < 0.0001), highlighting the advantage of the algal system in producing fully active, post-translationally modified hirudin. This superior activity is consistent with proper tyrosine sulfation occurring in the algal product, a modification absent in the prokaryotic and yeast-derived counterparts.

**Figure 2.**
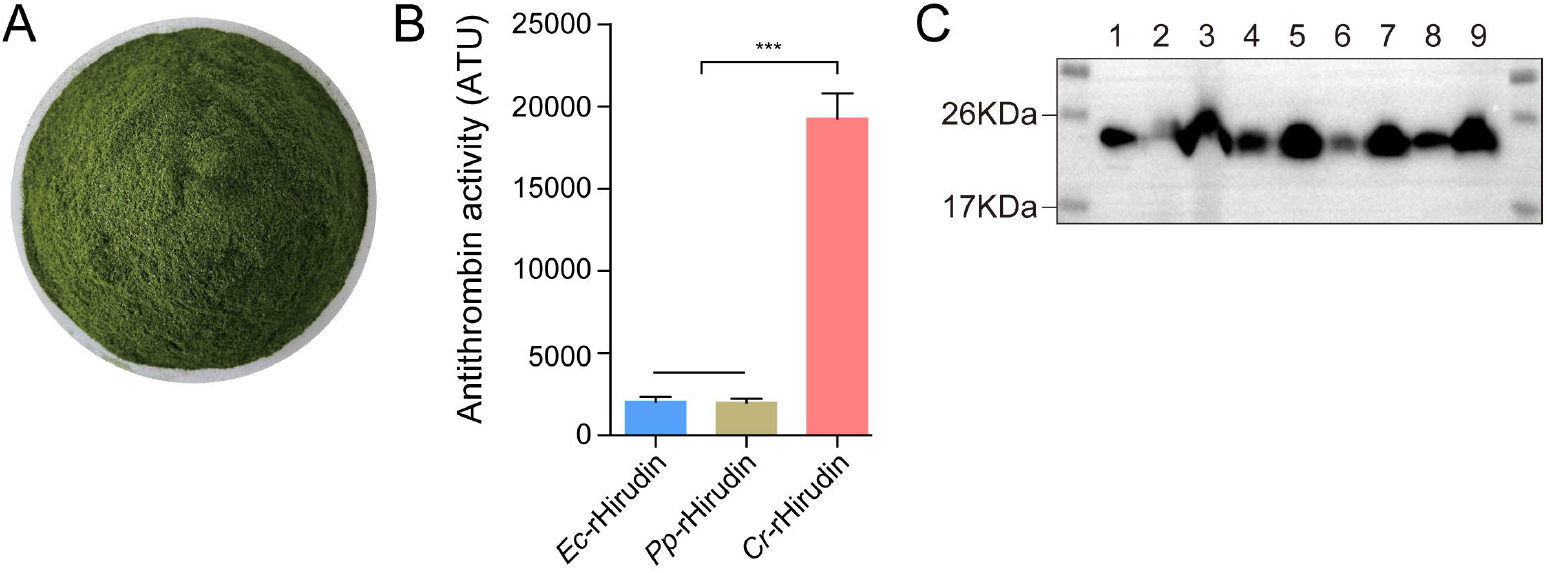
Lyophilized algal powder and activity of recombinant hirudin. (A) Photograph of the dry powdered *C. reinhardtii* biomass expressing hirudin. (B) Specific anticoagulant activity (ATU/mg) of recombinant hirudin produced in *E. coli* (*Ec*-rHirudin), *P. pastoris* (*Pp*-rHirudin), and *C. reinhardtii* (*Cr*-rHirudin). Data are presented as the mean ± SEM of three independent replicates. Statistical significance was determined by one-way ANOVA (***P < 0.0001). (C) Western blot analysis of HA-tagged hirudin stability in lyophilized biomass stored at room temperature over eight months. Samples were produced in either shake flasks or a bioreactor fermenter. The lanes represent algae powder produced under different conditions and storage durations: (1) on the same day by shake flask, (2) one month by fermentation, (3) one month by shake flask, (4) three months by fermentation, (5) three months by shake flask, (6) six months by fermentation, (7) six months by shake flask, (8) eight months by fermentation, (9) eight months by shake flask.

To assess the stability of the algal-produced hirudin, we pressed the lyophilized algal powder into tablet form and stored samples at room temperature for up to 8 months. Western blot analysis with an anti-HA antibody showed that hirudin protein levels remained detectable and stable throughout the storage period in all samples. Notably, this stability was observed in powders produced from both shake-flask cultures and 5 L fermenter batches, with no apparent degradation over time (Figure 2C). These results indicate that the recombinant hirudin in the algal powder maintains its integrity during long-term storage, which is advantageous for practical handling and formulation.

### *In vivo* anticoagulant efficacy in mice

An inferior vena cava (IVC) thrombosis model in mice was used to evaluate the anticoagulant efficacy of the hirudin-loaded algal powder when delivered orally. In untreated control mice (fed with wild-type algae powder lacking hirudin), large, dark-red thrombi developed in the IVC. By contrast, mice that received the recombinant hirudin algal powder had markedly reduced thrombus formation, with many exhibiting either very small, translucent clots or no visible thrombus at all (Figure 3A). Quantitative analysis of thrombus weight confirmed a significant reduction in the treated group: the average thrombus weight in hirudin-treated mice was dramatically lower than that of controls (***P < 0.0001). In the control group, 100% of mice (25/25) developed substantial clots, whereas only 20% (5/25) of the treated mice had any detectable thrombus, and those few clots were much lighter than control clots (Figure 3B). Approximately 80% of the treated animals showed complete prevention of thrombosis, indicating a potent antithrombotic effect of the orally administered hirudin-algal powder.

**Figure 3.**
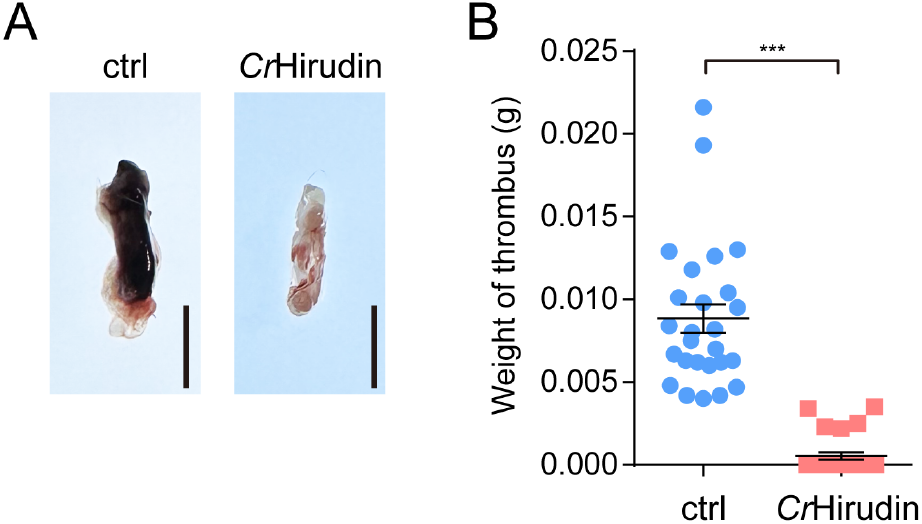
Oral administration of hirudin-expressing algae inhibits thrombosis in mice. (A) Representative images of thromboses in the IVC from a control mouse (left, ctrl) and a mouse treated with the hirudin algal powder (right, *Cr*Hirudin). Scale bar: 5mm. (B) Thrombus weights in individual mice from the control and treated groups. Each point represents the thrombus mass from one mouse, with mean ± SEM indicated. Mice that received oral *Cr*Hirudin show significantly lower thrombus weights compared to controls (***P < 0.0001). In fact, the majority of *Cr*Hirudin-treated mice had no visible thrombus (points at 0 mg), whereas all control mice developed sizable clots, underscoring the efficacy of the oral hirudin therapy.

### Safety assessment of oral algal hirudin

Potential anticoagulant-related side effects were evaluated by monitoring platelet counts and checking for bleeding in treated vs. control animals. Mice given the recombinant *C. reinhardtii*-hirudin powder maintained relatively stable platelet levels: all individuals exhibited <35% reduction in platelet count post-treatment, which is below the threshold for significant thrombocytopenia. In contrast, mice treated with the clinically used anticoagulants bivalirudin or heparin experienced substantial platelet reductions, with many showing >65% drops (Figure 4A). Thus, the incidence of thrombocytopenia was significantly lower in the *Cr*Hirudin-treated group compared to bivalirudin or heparin groups, indicating a reduced risk of heparin-induced thrombocytopenia-like effects.

**Figure 4.**
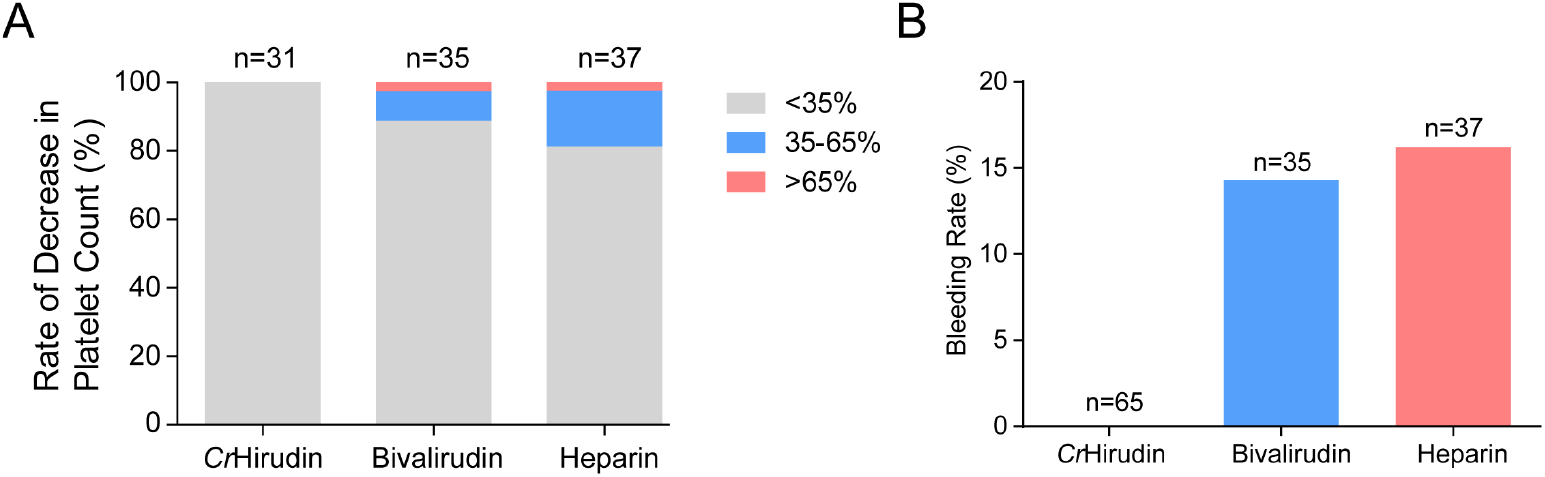
Comparative safety outcomes of algal hirudin and conventional anticoagulants in mice. (A) Incidence and severity of platelet count reduction after treatment. The stacked bars show the percentage of mice in each treatment group with mild (<35% drop), moderate (35–65% drop), or severe (>65% drop) thrombocytopenia. The number of mice per group (n) was 31 for *Cr*-Hirudin, 35 for bivalirudin, and 37 for heparin. (B) Percentage of mice exhibiting significant bleeding upon necropsy for each treatment group. The number of animals assessed (n) was 65 for *Cr*-Hirudin, 35 for bivalirudin, and 37 for heparin.

Furthermore, gross pathological examination at necropsy revealed no signs of hemorrhage in any of the mice that received the hirudin-algae oral treatment. There were no observable internal bleeds or subcutaneous hemorrhages in this group, suggesting that the fibrinolytic system was not excessively activated. In stark contrast, over 10% of the mice in the heparin and bivalirudin groups showed clear evidence of bleeding complications, such as extensive subcutaneous bruising or internal organ hemorrhages (Figure 4B). These results demonstrate that the algal-derived hirudin not only is effective at preventing thrombosis but also has a superior safety profile, with a markedly lower risk of bleeding and thrombocytopenic side effects compared to traditional anticoagulants.

## Discussion

Hirudin’s anticoagulant activity is critically dependent on its tertiary structure, stabilized by three disulfide bonds, and a sulfated tyrosine at position 63, both of which are essential for maximal activity^*33*^. Tyrosine sulfation, in particular, enhances the peptide’s affinity for thrombin’s active site, thereby dramatically increasing anticoagulant efficacy^*16,24*^. This tyrosine sulfation is catalyzed by tyrosylprotein sulfotransferase (TPST), an enzyme present in higher eukaryotes that selectively adds sulfate groups to specific tyrosine residues. Conventional recombinant expression platforms like *E. coli* and yeast, despite their low cost and high yield, lack TPST and other machinery for such post-translational modifications. Consequently, hirudin produced in these hosts remains unsulfated and exhibits only about 10% of the biological activity of native hirudin, creating a major bottleneck for industrial production^*34*^.

In this study, we addressed this limitation by leveraging *C. reinhardtii* as a eukaryotic expression chassis. Our results confirm that *C. reinhardtii*-derived hirudin undergoes tyrosine sulfation, which correlates with a dramatically enhanced anticoagulant potency. The algal-expressed hirudin achieved a specific activity of ∼20,000 ATU/mg, comparable to that of natural hirudin and far exceeding the activity of hirudin from *E. coli* or yeast (Figure 2B). This indicates that the *C. reinhardtii* expression environment can closely replicate the conditions required for proper folding and modification of complex peptides. While mass spectrometric analysis is needed to conclusively identify the exact sulfation site(s), the detection by an anti-sulfotyrosine antibody and the high functional potency strongly suggest that the critical Tyr-63 residue is correctly modified. Thus, the green alga-based system succeeds where traditional microbial systems fall short, yielding a fully active anticoagulant.

Another major finding of this work is the successful oral delivery of hirudin using a whole-cell microalgal formulation. Currently, hirudin and its analogs must be administered parenterally due to poor oral bioavailability, requiring frequent dosing and hospitalization that increase patient burden and infection risk^*17,35*^. In our murine thrombosis model, oral administration of lyophilized, hirudin-expressing *C. reinhardtii* powder effectively prevented thrombosis. This proof-of-concept for an oral, whole-cell biotherapeutic could revolutionize anticoagulant therapy by improving patient compliance and potentially providing a more sustained drug release profile. The robust cell wall of *C. reinhardtii*, a GRAS-certified organism, likely protects the peptide from degradation in the gastrointestinal tract and facilitates its uptake^*8*^.

Equally important, our oral hirudin platform demonstrated an excellent safety profile *in vivo*. Treated animals showed no signs of bleeding or thrombocytopenia, side effects commonly observed in those receiving injectable heparin or bivalirudin. The algal delivery system may mitigate these adverse effects by avoiding the high peak systemic concentrations that can cause bleeding or by eliminating animal-derived contaminants that can trigger immune reactions like heparin-induced thrombocytopenia. Overall, the ability to administer a safe and effective anticoagulant orally opens new avenues for the management of thromboembolic diseases, making therapy more patient-friendly and reducing the risks associated with injectable anticoagulants.

In conclusion, by harnessing the unique advantages of *C. reinhardtii* – its capacity for complex PTMs, its GRAS status, and its suitability for oral administration—we have established a green biomanufacturing platform that addresses key shortcomings of conventional hirudin production. This system produces hirudin with superior activity and an improved safety profile, potentially overcoming the challenges of short half-life and bleeding risk that plague current anticoagulant therapies. This study lays the groundwork for a next-generation oral anticoagulant. Future work will focus on detailed modification analysis, expression optimization, and industrial scale-up to translate this platform from bench to bedside.

## Notes

The authors declare no competing financial interest.

## Acknowledgments

This work has been financed by grants from the Natural Science Foundation for Distinguished Young Scholars of Hubei Province (2025AFA060), the Wuhan Municipal Education Bureau’s Program for the Integration of Research and Education (2025KCJ03), and the National Natural Science Foundation of China (32170702).

